# INF2-mediated actin polymerization at ER-organelle contacts regulates organelle size and movement

**DOI:** 10.1101/2024.07.06.602365

**Authors:** Cara R. Schiavon, Yuning Wang, Jasmine W. Feng, Stephanie Garrett, Tsung-Chang Sung, Yelena Dayn, Chunxin Wang, Richard J. Youle, Omar A. Quintero-Carmona, Gerald S. Shadel, Uri Manor

**Affiliations:** Department of Cell & Developmental Biology, University of California, San Diego; Department Of Biology, University of Richmond, VA; Salk Institute for Biological Studies; National Institute of Neurological Disorders and Stroke

## Abstract

Proper regulation of organelle dynamics is critical for cellular function, but the mechanisms coordinating multiple organelles remain poorly understood. Here we show that actin polymerization mediated by the endoplasmic reticulum (ER)-anchored formin INF2 acts as a master regulator of organelle morphology and movement. Using high-resolution imaging, we demonstrate that INF2-polymerized actin filaments assemble at ER contact sites on mitochondria, endosomes, and lysosomes just prior to their fission. Genetic manipulation of INF2 activity alters the size, shape and motility of all three organelles. Our findings reveal a conserved mechanism by which the ER uses actin polymerization to control diverse organelles, with implications for understanding organelle dysfunction in neurodegenerative and other diseases. This work establishes INF2-mediated actin assembly as a central coordinator of organelle dynamics and inter-organelle communication.

## Results and Discussion

Alterations in organelle dynamics, such as organelle fission, mobility, and changes in inter-organelle contacts, are implicated in multiple human diseases, particularly various neuropathies^1-7^. The ER (endoplasmic reticulum) makes extensive contacts with organelles throughout the cell. Extensively-studied ER-mitochondria contacts play roles in important cellular processes, including organelle dynamics^8, 9^. ER tubules mark mitochondrial fission sites by wrapping and constricting mitochondria until division^10, 11^. Additional reports have demonstrated ER tubules marking fission events of other organelles as well^12, 13^.

Organelle dynamics are also regulated by the cytoskeleton. Nearly all organelles move throughout the cell along microtubules and/or actin filaments. Actin also plays a key role in regulating mitochondrial dynamics. Prior to fission, actin accumulates on mitochondria which provides the force necessary to constrict the mitochondrion and ultimately cause fission^14-16^. Proper coordination of this process is essential for overall cellular health, as mitochondrial dynamics directly impact important functions including lipid synthesis, energy production, calcium signaling, and mitochondrial turnover. Like mitochondria, organelles of the endolysosomal pathway are highly dynamic and their size, fission, fusion, and positioning must remain balanced to maintain proper function^17, 18^. However, the regulation of endolysosome dynamics, particularly the involvement of inter-organelle contacts and actin, has so far not been extensively studied.

Given that organelle fission has been associated with both the ER and the actin cytoskeleton, we hypothesized that ER-associated actin (“ER-actin”) may be responsible for organelle fission. To test this hypothesis, U2OS cells were co-transfected with plasmids directing expression of a fluorescently-tagged organelle marker and an ER-targeted actin nanobody previously shown to label ER-associated actin (“AC-ER”)^19^. Prior work showed that this probe accumulates at ER-mitochondria contacts during mitochondrial fission events^19^, and we hypothesized that we should be able to observe similar results for other organelles if ER-actin is involved in their fission. Following transfection, cells were imaged via time-lapse Airyscan confocal microscopy. Fission events in the time-lapses were manually identified in a blinded fashion by visualizing the organelle marker alone, then later scored for the presence or absence of AC-ER enrichment (**Fig. 1**). To image endosomes, cells were co-transfected with the early endosome marker Rab5-mCherry (magenta) and AC-ER GFP (green). Lysosomes were imaged by transfecting LAMP1-mCherry, and mitochondria were imaged with the MitoTracker Deep Red dye. Enrichment of the AC-ER signal was evident in 96-100% of the 70 fission events scored, far greater than the amount expected by chance. This enrichment was evident during organelle constriction prior to fission, consistent with the model that actin promotes organelle constriction and eventual fission.

**Figure 1:**
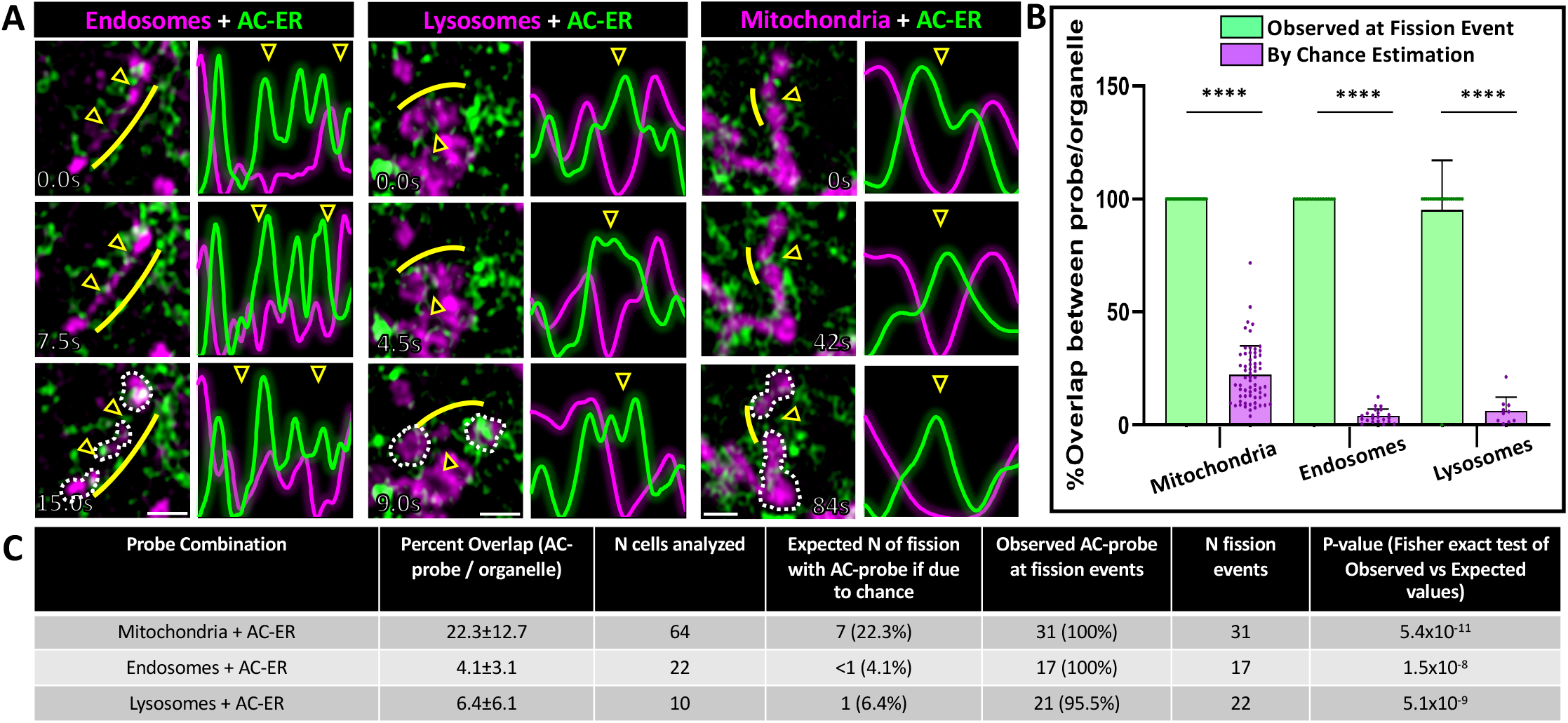
ER-associated actin accumulates at endosomal, lysosomal, and mitochondrial fission sites prior to organelle division. U2OS cells were co-transfected with plasmids directing expression of fluorescently-tagged organelle markers and fluorescently-tagged, ER-targeted actin nanobodies (“AC-ER”) and timelapse videos were collected. **A**. Endosomes are labeled with Rab5-mCherry, lysosomes were labeled with LAMP1-mCherry, and mitochondria were labeled with MitoTracker Deep Red (magenta). AC-ER GFP is shown in green. Example organelles are shown at different time points prior to and immediately after the fission event. Arrowheads indicate the fission sites. White dotted lines have been drawn around the “daughter organelles” resulting from the fission event in the bottom panel for clarity. To aid visualization, line scans were drawn over the fission sites and the surrounding regions. The resulting pixel intensities associated with the line scans are shown. Yellow lines indicate the region where the line scan was drawn (shifted so as not to block visualization of the organelle). Scale bars are 1µm. **B**. Graph comparing the frequency of observed presence/absence of AC-ER at organelle fission events (green bars) versus the possibility of AC-ER being at fission events by chance (purple bars). Actual observed AC-ER at fission events was determined by manually scoring fission events in a blinded fashion, followed by looking for AC-ER signal at the identified fission events. By chance values were determined by calculating the percent of organelle signal overlapped by AC-ER signal (described in greater detail Supp. Fig 1D and Methods). Observed versus by chance values were statistically compared via Fisher’s exact test. **C**. Tabular breakdown of the n-values and p-values associated with AC-ER fission event analysis. For each organelle + AC-ER combination the n-values corresponding to the number of cells analyzed as well as the number of fission events scored are displayed. The calculated percent overlap (corresponding to the purple bars in B) are shown in the second column. These values were used to calculate the “expected” percentage of fission events where AC-ER would be present if due to chance. The observed and expected percentages were compared via Fisher’s exact test and the resulting p-values are displayed in the last column (corresponding to the p-values in B). All experiments were performed with N=3 biological replicates.

Although AC-ER hotspots are associated with ER-actin, the AC-ER probe is targeted to the entirety of the ER membrane. Thus, there is a possibility that AC-ER signal may be present at an organelle fission event simply due to the extensive overlap between the ER and other organelles in the cell. To rule out this possibility, masks of the organelle and AC-ER channels were overlaid to determine the percentage of overlap between the organelle and AC-ER. The fraction of the organelle overlapped by AC-ER is our estimation of the probability that AC-ER is present at a fission event “by chance” (purple bars in the graph, **Fig. 1B, Supp. Fig. 1D**). We also checked every identified organelle fission event for the presence or absence of thresholded AC-ER signal (green bars in the graph). AC-ER was present at nearly 100% of fission events, significantly higher than the values calculated by chance. Thus, the observed presence of AC-ER at organelle fission events is unlikely to be coincidental. **Fig. 1C** shows the tabular breakdown of the n-values and p-values associated with AC-ER fission event analysis.

To distinguish general ER from AC-ER accumulation at fission events, cells were co-transfected with BFP-KDEL as a general ER marker. BFP-KDEL did not display the same degree of accumulation as AC-ER at fission sites (**Supp. Fig. 1A-C, Supp. Videos 1-3**). Additionally, mitotracker-labeled cells were also co-transfected with DRP1-mCherry, which labels mitochondrial fission sites. We observed DRP1-mCherry and AC-ER at each mitochondrial fission site, providing further evidence that the events being scored are true mitochondrial fission events. Together, our results suggest a general cellular mechanism by which ER-actin promotes the fission of not just mitochondria, but also lysosomes and endosomes.

It has been shown that inhibition of actin polymerization leads to reduced mitochondrial fission events and increased mitochondrial elongation^15, 20-22^. Given that ER-actin is consistently present at other organelle fission sites, we posited that actin is playing a role in the regulation of organelle fission and, more broadly, organelle shape. We thus hypothesized that a loss of actin would directly impact organelle fission rates and, as a result, organelle morphology. To test this hypothesis, we transfected U2OS cells with fluorescent organelle markers and imaged the cells live following addition of Latrunculin B or vehicle control. Latrunculin B (LatB) leads to a dramatic loss of filamentous actin throughout the cell. Because this is acutely toxic to cells, a low dose (200nM) was used to allow cells to survive long enough (60-90 minutes) to observe changes in organelle morphology.

Consistent with previous reports, we observed that mitochondria appear elongated following LatB treatment. In support of our hypothesis of a more general mechanism, endosomes and lysosomes also appeared larger in size and trended towards a less spherical shape following treatment with LatB (**Supp. Fig 2**). Quantification revealed a significant increase (∼80%) in the area of individual organelles following treatment with LatB for all three organelles tested (**Supp. Fig 2B)**. Because we hypothesized that the increase in organelle size is a result of a change in the organellar fission rate, we manually scored the number of fission events occurring in timelapses following vehicle control or LatB treatment. Scoring of fission events revealed a significant decrease (40-52%, **Supp. Fig 2C**) in both endosomal and lysosomal fission rates following LatB treatment. Together, our results are consistent with a model in which actin filaments play a key role in promoting organelle fission.

**Figure 2:**
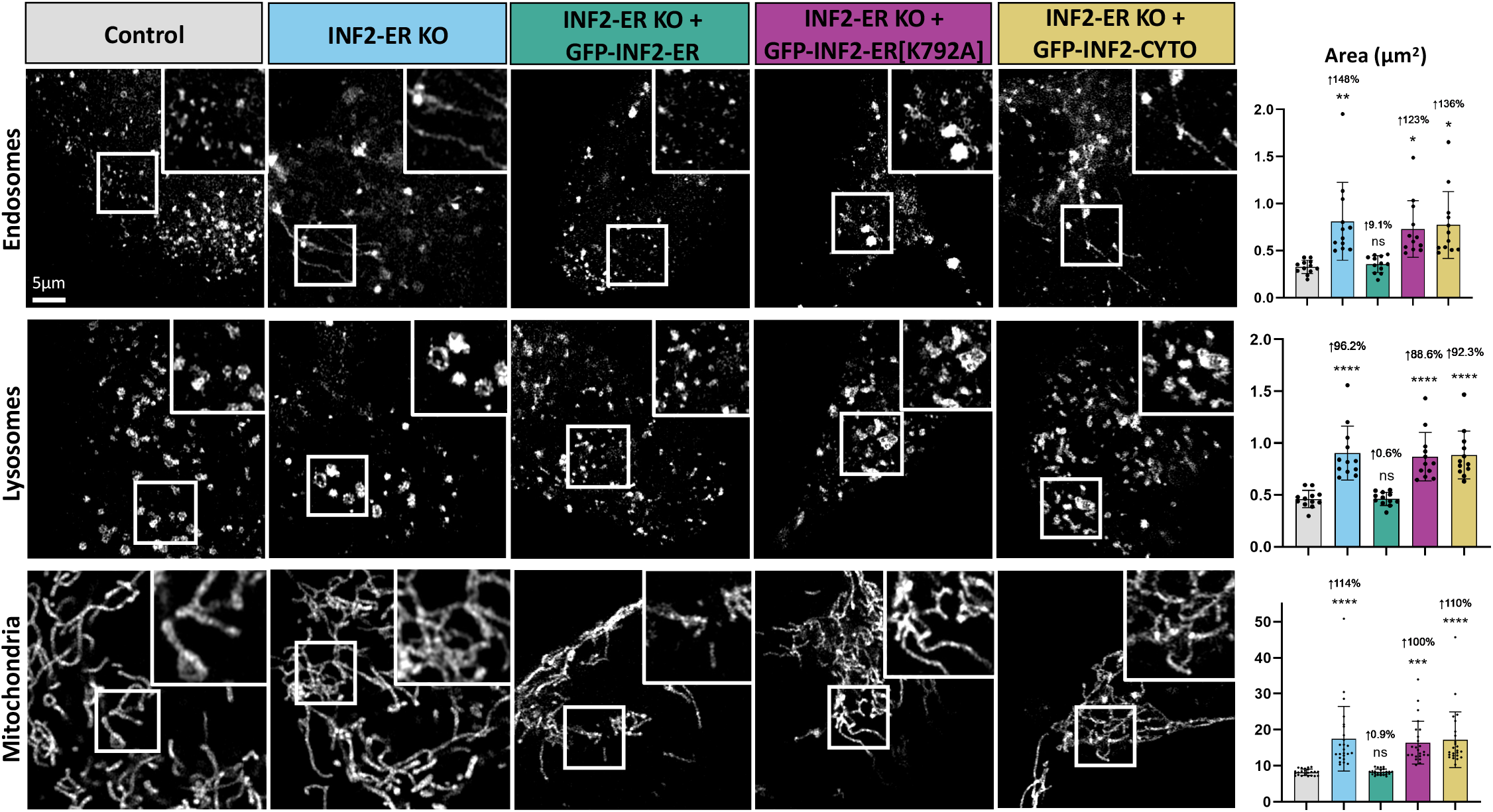
Loss of INF2 causes organelle enlargement and elongation. Experiments were carried out in wild-type U2OS cells (“Control”) and U2OS cells where the ER-anchored isoform of INF2 has been knocked-out (“INF2-ER KO”). Cells were imaged live following transfection with Rab5-mCherry to label endosomes, LAMP1-mCherry to label lysosomes, or staining with MitoTracker Deep Red to label mitochondria. The leftmost column displays representative images of each organelle in Control U2OS cells. The following column shows each organelle in INF2-ER KO cells. The following column displays INF2-ER KO cells that have been co-transfected with INF2-ER GFP. The following column displays INF2-ER KO cells co-transfected with INF2-ER[K792A] GFP. The following column displays INF2-ER KO cells co-transfected with INF2-Cyto GFP. The last column displays quantification of individual organelle area for each condition. Black dots show the average per cell and bars indicate the average per condition. Error bars show standard deviation. N = 12 cells per condition for endosomes and lysosomes and 24 cells per condition for mitochondria. For all graphs, the magnitude of the change and p-value compared to control is shown. An up arrow indicates an increase and a down arrow indicates a decrease. For statistical comparisons ^****^ indicates p-value≤0.0001, ^***^ indicates p-values≤0.001, ^**^ indicates p-value≤0.01, ^*^ indicates p-value≤0.05, and ns indicates p-value>0.05. Conditions were compared via ordinary one-way ANOVA. All experiments were performed with N=3 biological replicates.

Because there are various factors that may increase mitochondrial area (i.e. length, interconnectivity), we further investigated mitochondrial morphology. We found that mitochondrial length is significantly increased (27%, **Supp. Fig 2D**) following LatB treatment, while the number of branches per mitochondrion is decreased (34%, **Supp. Fig 2D**) after normalizing to mitochondrial length. Thus, the increase in mitochondrial area is primarily driven by an increase in mitochondrial length. In addition to the increased area of endosomes and lysosomes, a significant decrease (9-12%, **Supp. Fig 2E**) in circularity for both endosomes and lysosomes was observed following LatB treatment. Together, these results support the hypothesis that actin filaments are important for regulating organelle morphology.

Given that ER-actin appears to promote organellar fission, we pursued the ER-anchored splice isoform of INF2 (Inverted Formin 2) as a potential regulator of this process. INF2 is a formin protein with two splice isoforms which localize to either the cytosol (“INF2-CYTO”) or the ER (“INF2-ER”). Furthermore, INF2-ER has previously been shown to play a role in the promotion of mitochondrial fission via its actin polymerizing activity^15^. Experiments were carried out in wild-type U2OS cells (“Control”), U2OS cells where INF2-ER has been knocked-out (“INF2-ER KO”), and U2OS cells where all isoforms of INF2 have been knocked-out (“Complete INF2 KO”). Cells were imaged live following transfection with either Rab5-mCherry to label endosomes or LAMP1-mCherry to label lysosomes, or staining with MitoTracker Deep Red to label mitochondria. We observed cells lacking INF2-ER had elongated mitochondria, as previously reported (**Fig. 2**). We also observed that endosomes in INF2-ER KO cells were also significantly enlarged and often tubular in shape. INF2-ER KO lysosomes were also significantly enlarged and appeared less spherical than their WT counterparts.

To control for off-target effects, INF2-ER KO cells were rescued via exogenous expression of GFP-INF2-ER. In this condition, endosomes, lysosomes, and mitochondria were restored to a morphology indistinguishable from WT cells. This indicates that the organelle morphology phenotypes observed in INF2-ER KO cells was indeed a result of loss of INF2-ER rather than an off-target effect. To test the hypothesis that INF2-ER impacts organelle morphology via its actin polymerizing activity, we transfected INF2-ER KO cells with GFP-INF2-ER[K792A]. K792A is a point mutation that disrupts INF2’s ability to effectively polymerize actin^23^. Expression of GFP-INF2-ER[K792A] did not reverse the phenotype observed in INF2-ER KO cells, supporting the hypothesis that INF2-mediated regulation of organelle shape is dependent on its actin polymerizing activity.

We also compared the contributions of INF2-CYTO vs INF2-ER. To this end, we attempted rescue by exogenous expression of GFP-INF2-CYTO. Expression of GFP-INF2-CYTO also did not reverse the phenotype observed in INF2-ER KO cells, which supports the model that the regulation of organelle shape is predominantly modulated by the ER-anchored isoform of INF2.

Quantification of organelle area supports the conclusion that loss of INF2-ER causes organelle enlargement and elongation. INF2-ER KO cells showed an average increase of 96-148% (**Fig. 2**) in individual organelle area compared to control. This increase is largely maintained in both GFP-INF2-ER[K792A] and GFP-INF2-CYTO rescue conditions. In contrast, the GFP-INF2-ER rescue condition resulted in organelles with an area very similar to control with a maximum increase of only 9% (**Fig. 2**). In addition to area, individual mitochondrial length and the number of branches per mitochondria were also quantified (**Supp. Fig 3**). Similar to the results with LatB, mitochondrial length was significantly increased (62% **Supp. Fig 3**) in INF2-ER KO cells, but branching was not increased. The circularity of endosomes and lysosomes was also measured for all conditions. In agreement with the observed tubulation phenotypes, endosome circularity was significantly reduced (20%, **Supp. Fig 3**) in INF2-ER KO conditions. Lysosome circularity was also reduced to a more modest but still statistically significant degree (8%, **Supp. Fig 3**). These alterations were reversed upon expression of GFP-INF2-ER but not GFP-INF2-ER[K792A] or GFP-INF2-CYTO expression.

**Figure 3:**
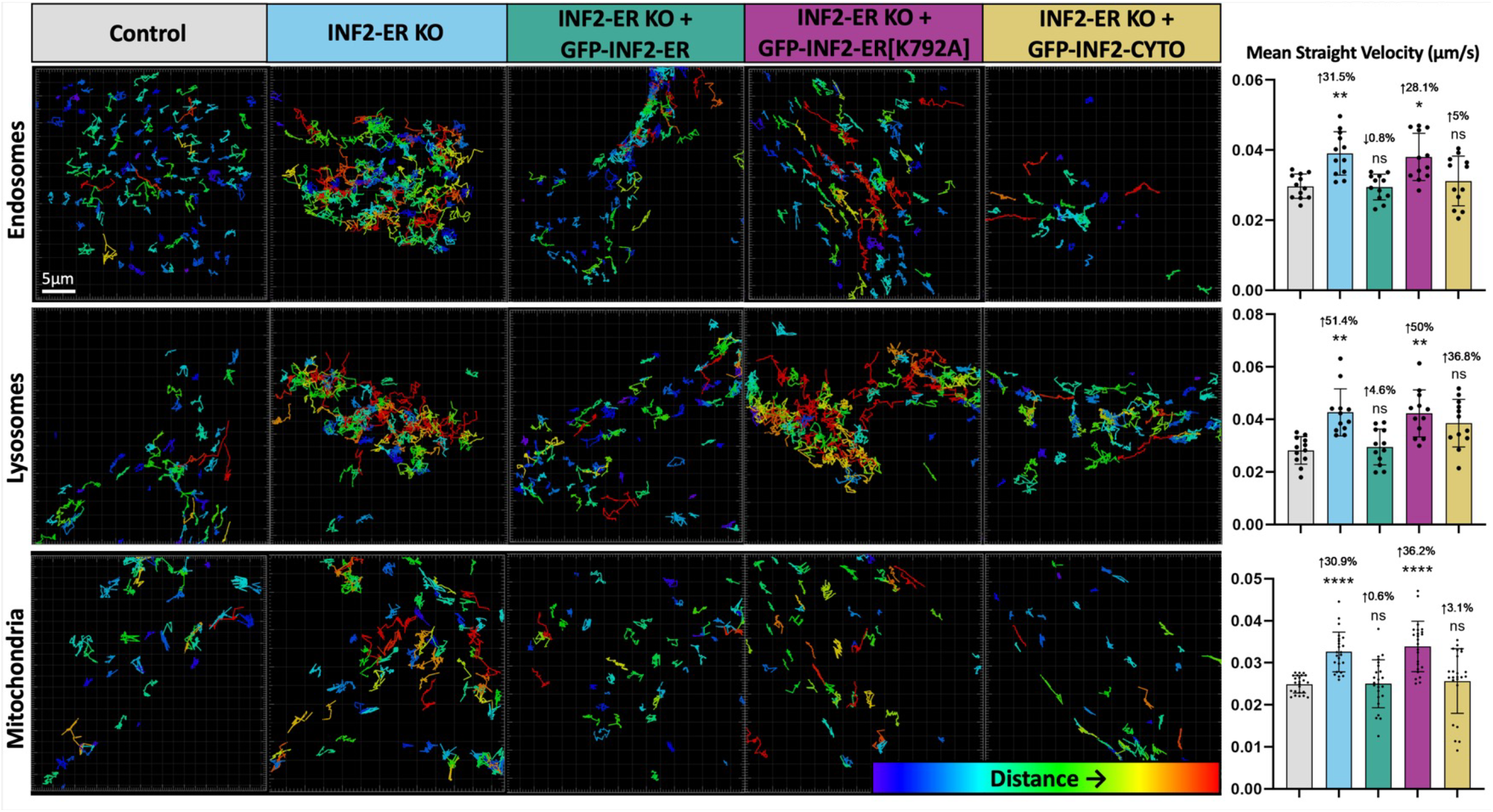
Loss of INF2 increases organelle mobility. Organelles were labeled in wild-type U2OS cells (control) and INF2-ER KO U2OS cells using Rab5-mCherry (endosomes), LAMP1-mCherry (lysosomes), or MitoTracker Deep Red (mitochondria) and their movement was tracked during 5-minute timelapses using Imaris software. To aid in visualization of organelle movement, tracks marking the trajectories of each organelle in representative cells are shown. These tracks are color-coded according to Imaris’ calculated “track displacement distance”. This is the measured distance between the organelle’s location at the beginning and its location at the end of the timelapse. Thus, more red tracks indicate organelles that cover more distance, and therefore, move more, while more blue tracks indicate organelles that move less. The first column shows tracks generated from endosome, lysosome, and mitochondrial movement in the Control condition. The following column displays organelle tracks from the INF2-ER KO condition. The following column displays tracks from INF2-ER KO cells co-transfected with INF2-ER GFP. The following column displays tracks from INF2-ER KO cells co-transfected with INF2-ER[K792A] GFP. The following column shows tracks from INF2-ER KO cells co-transfected with INF2-Cyto GFP. The last column shows quantification of organelle movement based on Imaris organelle tracking. To correct for tracks that do not last the entire duration of the timelapse due to organelles leaving/entering the frame or going in/out of focus, the “Mean Straight Velocity” of each organelle was calculated by dividing the track displacement distance by the track duration. N = 12 cells per condition for endosomes and lysosomes and 24 cells per condition for mitochondria. For all graphs, black dots show the average per cell and bars indicate the average per condition. Error bars show standard deviation. The magnitude of the change and p-value compared to control is shown. An up arrow indicates an increase and a down arrow indicates a decrease. For statistical comparisons ^****^ indicates p-value≤0.0001, ^***^ indicates p-values≤0.001, ^**^ indicates p-value≤0.01, ^*^ indicates p-value≤0.05, and ns indicates p-value>0.05. Conditions were compared via ordinary one-way ANOVA. All experiments were performed with N=3 biological replicates.

Because we suspect that the alterations in organelle morphology are a result of changes in fission rate in the INF2-ER KO conditions, endosome and lysosome fission rates were also measured. Although results were not statistically significant compared to control, there was a trend towards a reduction in fission rate (41-43%, **Supp. Fig 3**) in the INF2-ER KO line which was reversed upon expression of GFP-INF2-ER but not GFP-INF2-ER[K792A] for both endosomes and lysosomes.

All the imaging experiments and quantification described thus far for INF2-ER KO cells were also conducted using Complete INF2 KO cells in which neither the cytosolic or ER-anchored splice isoform of INF2 was present. When comparing equivalent INF2-ER KO and Complete INF2 KO conditions, we observed nearly identical KO and rescue phenotypes (**Supp. Fig. 3**). This further supports the model that modulation of organelle fission rate and morphology is primarily regulated by INF2-ER, as exogenous expression of GFP-INF2-ER resulted in an almost complete reversal of the organelle phenotypes in Complete INF2 KO cells, while expression of GFP-INF2-CYTO failed to rescue a WT phenotype.

Given that loss of INF2 causes a dramatic change in organelle morphology, we sought to test the effect(s) of increased INF2 activity on organelle morphologies. The A149D point mutation prevents the autoinhibition of INF2, resulting in an excess of actin polymerizing activity and mitochondrial fragmentation^15^. In WT U2OS cells transfected with GFP-INF2-ER[A149D], mitochondria were noticeably fragmented, in agreement with previously published results^15^ (**Supp. Fig 4A-B**). This is unlikely to be an artifact of overexpression as transfection of wild-type GFP-INF2-ER did not significantly alter mitochondrial morphology. Mitochondrial fragmentation was not observed in GFP-INF2-ER[A149D] expressing cells after 15-30min treatment with 1µM LatB, indicating that the mitochondrial fragmentation observed was actin-dependent. In agreement with the observed mitochondrial fragmentation, mitochondrial quantification revealed significantly shorter (47%, **Supp. Fig 4C**) mitochondria following expression of GFP-INF2-ER[A149D] but not wild-type GFP-INF2-ER. Treatment with LatB restored and even increased (26%, **Supp. Fig 4C**) mitochondrial length, similar to what was observed in WT cells treated with LatB. Mitochondrial branching was unchanged following expression of wild-type GFP-INF2-ER and decreased (45%, **Supp. Fig 4C**) when normalizing to mitochondrial length in the LatB condition, also similar to WT cells treated with LatB. Expression of GFP-INF2-ER[A149D] caused a reduction in mitochondrial branching (26%, **Supp. Fig 4C**), even after normalizing to the shorter mitochondrial length.

Unlike mitochondria, expression of GFP-INF2-ER[A149D] did not significantly reduce endosome or lysosome size, possibly because these organelles rarely tubulate during normal conditions and thus have already reached a “floor” in fission-induced size reduction. The GFP-INF2-ER[A149D] + LatB condition resulted in enlargement (55-78%, **Supp. Fig. 4B**) of both organelles, similar to the results obtained in WT cells treated with LatB (**Supp. Fig. 2**). Similarly, expression of GFP-INF2-ER[A149D] did not further increase the circularity of endosomes or lysosomes, though when combined with LatB treatment we did observe reduced endosome and lysosome circularity (7-11%, **Supp. Fig 4D**), similar to results in WT cells treated with LatB (**Supp. Fig. 2**).

While imaging organelles in live INF2 KO cells, we noted apparent differences in their movement when compared to control cells. To further investigate this, we labeled organelles as described previously in wild-type, INF2-ER KO, and Complete INF2 KO U2OS cells and tracked the movement of organelles during 5-minute time-lapse windows. Organelle tracking was performed with commercial Imaris software. To aid in visualization of organelle movement, tracks marking the trajectories of each organelle in representative cells are shown in **Fig. 3**. Videos corresponding to these tracks are in **Supp. Videos 4-6**. These tracks are color-coded according to the measured distance between the organelle’s location at the beginning and end of the time-lapse. Thus, more red tracks indicate organelles that cover more distance, and therefore, move more, while more blue tracks indicate organelles that move less. In the INF2 KO conditions, endosomes, lysosomes, and mitochondria all move more compared to control as indicated by longer, higher displacement distance tracks. In INF2 KO + GFP-INF2-ER conditions, organelle movement was similar to control, indicating that, similar to the morphological phenotypes (**Fig. 2**), the organelle mobility phenotype was not due to an off-target effect. Tracks from the INF2 KO + GFP-INF2-ER[K792A] conditions were similar to those from the INF2 KO conditions, indicating INF2’s actin polymerizing activity was necessary to mediate its regulation of organelle movement. In the INF2 KO + GFP-INF2-CYTO conditions, track displacement appeared intermediate between the tracks typically observed in control versus INF2 KO conditions. This indicates that, although the ER-anchored isoform of INF2 clearly contributes to regulation of organelle mobility, overexpressing the cytosolic isoform may also have an impact.

To account for tracks that do not last the entire duration of the time-lapse due to organelles leaving/entering the frame or going in/out of focus, the “Mean Straight Velocity” of each organelle was also calculated by dividing the track displacement distance by the track duration. Quantification reveals values that are consistent with the observed tracks for each condition, showing a 31-51% increase in organelle mean straight velocity in the INF2-KO conditions compared to control (**Fig 3, Supp. Fig 5**). For expression of GFP-INF2-CYTO, the mean value was not statistically different from control, but there remains a slight trend towards increased organelle movement (3-37% **Fig. 3, Supp. Fig 5**), particularly for lysosomes. This is consistent with INF2-CYTO rescuing the organelle mobility phenotype though not to the same degree as INF2-ER.

In addition to the Imaris quantification, organelle movement was measured via a complementary “autocorrelation” method which measures the correlation coefficient when comparing the first frame of the time-lapse to all consecutive frames of the same timelapse (**Supp. Fig 5**)^24^. As the time-lapse progressed and organelles moved, the correlation when comparing to the first frame decreases at a rate inversely proportional to the rate of organelle movement. When applying this method to the timelapse data, a drop in the correlation coefficient over time was observed for all conditions but was significantly more pronounced for the INF2 KO as well as the INF2 KO + GFP-INF2-ER[K792A] conditions. This agrees with the Imaris organelle tracking results showing a significant increase in organelle movement in these conditions compared to control. Also similar to the object tracking results, the INF2 KO + GFP-INF2-ER conditions generated curves nearly identical to the control curve, further indicating that restoration of the ER-anchored isoform of INF2 was sufficient to completely reverse the organelle mobility phenotype. Correlation values for the INF2 KO + GFP-INF2-CYTO conditions were more similar to the control curve than the INF2 KO curves but not identical to the control curve, suggesting that expression of the cytosolic isoform of INF2 partially rescued the organelle mobility phenotype.

Because INF2’s modulation of organelle movement depends on its actin polymerizing activity, we tested whether generalized inhibition of actin polymerization would alter organelle movement by treating wild-type U2OS cells with LatB (**Supp. Fig 6, Supp. Video 7**). Cells were transfected to label organelles as described previously, treated with 200nM LatB or vehicle control for 60-90 minutes, then imaged over time for an additional 5 minutes. Organelle movement was measured using both Imaris object tracking and the autocorrelation method. Both object tracking and autocorrelation results indicated a significant increase (33-52%, **Supp. Fig 6**) in organelle mobility following LatB treatment, suggesting that actin not only modulates organelle morphology but also organelle movement. This is consistent with the model that reduction of filamentous actin results in less hindrance of organellar movement.

Given the effect of INF2 KO on organelle mobility, we tested the effect of dominant active INF2 expression on organelle movement. These experiments were also motivated by previously published work showing that expression of INF2[A149D] caused a marked reduction in mitochondrial mobility^15^. Organelles were labeled as previously described and co-transfected with GFP (control), wild-type GFP-INF2-ER, or GFP-INF2-ER[A149D]. The GFP-INF2-ER[A149D] condition was also compared with and without 15-30 minute treatment with 1µM LatB (**Supp. Fig 7, Supp. Videos 8-10**). While control and wild-type GFP-INF2-ER tracks were similar, tracks from the GFP-INF2-ER[A149D] condition were noticeably shorter, indicating a reduction in organelle movement following expression of GFP-INF2-ER[A149D] but not wild-type GFP-INF2-ER. Thus, the reduction in organelle mobility is unlikely to be an artifact of overexpression but rather an effect specifically caused by expression of the dominant active mutant. Importantly, although GFP-INF2-ER[A149D] expression had differential results on organelle morphology depending on the organelle analyzed (only altering mitochondrial size), it consistently caused reduction of organelle movement (23-41%, **Supp. Fig 7**) for all three organelles studied. Conversely, tracks were longer (32-49%, **Supp. Fig 7**) in the GFP-INF2-ER[A149D] + LatB condition, consistent with LatB causing an increase in organelle movement and the decrease in organelle movement being actin-dependent. Autocorrelation analysis was consistent with the object tracking results, showing a faster decrease in autocorrelation for the GFP-INF2-ER[A149D] + LatB condition and a slower decrease for the GFP-INF2-ER[A149D] condition compared to control.

Taking all these results into account, we posit a model in which ER-actin accumulates on organelles, resulting in their constriction and eventual fission. The force necessary to constrict the organelles may be generated by actin polymerization, actin-associated myosin motor proteins, or other potential mechanisms. In addition to force generation, actin may also play an important role in recruiting and stimulating additional fission factors, such as dynamin-family proteins^14, 25, 26^. It is important to note that ER-actin is transient and represents a very small fraction of the overall cellular actin, so it may be difficult or even impossible to clearly observe its localization using traditional actin labeling methods as opposed to the ER-targeted actin chromobody AC-ER probes used in this study.

Analogous to its role in promoting mitochondrial fission, INF2 also promotes endosome and lysosome fission via its actin polymerizing activity. Regulation of organelle fission is mediated by the ER-anchored isoform of INF2 but not the cytosolic isoform. One possibility is that the close interaction between the ER and other organelles at precise locations prior to their fission allows the ER-anchored INF2 to be properly positioned to mediate actin polymerization, which promotes organelle constriction. Actin also facilitates recruitment of DRP1 (for mitochondria) and possibly other dynamin-related proteins to assemble and complete organelle fission. Of note, loss of INF2 reduces organelle fission rates but does not completely abolish fission. Thus, there are likely other actin-regulatory proteins involved in this process, an important topic for future studies.

We also propose that INF2’s actin polymerizing activity regulates organelle movement since organelle displacement is impacted both by loss of INF2 (increased movement) or expression of dominant active INF2 (decreased movement). In this instance, both INF2 isoforms contribute, but the ER-anchored isoform exerts the strongest influence. There are various mechanisms by which ER-actin accumulation may impede organelle movement including steric hindrance, alterations in availability of motor protein binding sites, alterations in microtubule post-translational modifications, or changes to other proteins that regulate organelle mobility, though this would require further study to determine. Together, our findings position INF2 and ER-actin at ER-organelle contacts as important regulatory factors for both organelle morphology and transport, key features that must be dynamically modulated to maintain proper organelle and cellular function. We speculate these effects may be important for the pathophysiology of Charcot-Marie-Tooth disease mutations in INF2^27^.

## Materials and Methods

### Cell culture, transfection, labeling, and drug treatments

U2OS cells were purchased from the American Type Culture Collection (ATCC Cat. # HTB-96). Complete INF2 KO U2OS cells were a generous gift from the Higgs lab and are described in Chakrabarti et al.^28^. INF2-ER KO U2OS cells were generated via CRISPR-mediated knock-out of the ER-anchored splice isoform of INF2. To generate INF2-ER KO U2OS cells, two gRNAs were used to delete exon 22: 5’ gRNA: GCAGCCACTTGCCTGGGACC and 3’ gRNA: TGGGGGCTAACAGCAGCTGC. gRNA sequences were cloned into gRNA_Cloning vector (Addgene plasmid #41824, a gift from George Church) and co-transfected with pEYFP-C1 into U2OS cells. GFP-positive cells were FACS sorted and plated into 96-well plates. Single colonies were screened with PCR using the following primer set (Fwd: GGAGAGGTGACTTGGGTGCG, Rev: GACACCAGACAGGAGCAACC). WT clones produce a 757 bp band whereas KO clones produce a band 257 bp shorter^29^. Cells were maintained in DMEM supplemented with 10% FBS at 37°C with 5% CO_2_. Cells were transfected using Lipofectamine 2000 (Thermo Fisher) according to the manufacturer’s instructions. Cells were plated onto 8-well no.1.5 imaging chambers (Cellvis) that were coated with 10 μg/mL fibronectin in PBS at 37°C for 30 min before plating. MitoTracker Deep Red (50 nM; Thermo Fisher) was added to cells for 30 min, and then cells were washed and allowed to recover for at least 30 min before imaging in FluoroBrite (Thermo Fisher) medium. After plating, for some experiments, cells were treated with Latrunculin B (Abcam) at a final concentration of 1µM for 15-30min or 200nM for 60-90min prior to imaging. Control conditions from the same experiments were treated with an equal volume of ethanol as a vehicle control.

### Airyscan confocal imaging

Cells were imaged with a Plan-Apochromat ×63/1.4 NA oil objective on an inverted Zeiss 880 LSM Airyscan confocal microscope with the environmental control system supplying 37°C, 5% CO_2_ and humidity for live-cell imaging. The GFP channels were imaged with a 488-nm laser line at ∼50-µW laser power. The mCherry channels were imaged with a 561-nm laser at ∼200-μW laser power. The MitoTracker Deep Red channel was imaged with ∼20-µW laser power. The BFP channel was imaged with a 405-nm laser line at ∼10-µW laser power. For time-lapse imaging, the zoom factor was set between 3× and 6× to increase the frame rate. In all cases, the maximum pixel-dwell time (∼0.684 μs per pixel) and 2× Nyquist optimal pixel size (∼40 nm per pixel) were used.

### Image processing and analysis

After acquisition, images were Airyscan processed using the auto-filter 2D-SR settings in Zen Blue (ZEISS). All images were post-processed and analyzed using Imaris (BITPLANE) and Fiji software.

### Plasmids

Rab5-mCherry (#27679), LAMP1-mCherry (#55073), DRP1-mCherry (#49152), and BFP-KDEL (#49150) plasmids were obtained from Addgene. AC-ER GFP was generated starting from the commercial vector of AC-tagGFP (ChromoTek) and cloned via the BglII and NotI restriction sites. The following amino acid sequence, derived from the C-terminal membrane binding region of cytochrome b5, was attached to the C-terminal portion of the AC probe to target the protein to the ER: IDSSSSWWTNWVIPAISAVAVALMYRLYMAED. The AC-ER GFP construct is described in more detail in Schiavon et al.^19^. The original INF2-ER GFP, INF2-ER[A149D] GFP, and INF2-CYTO GFP plasmids were a generous gift from the Higgs laboratory^15, 28^. All INF2 ORFs were cloned into the lentiviral pSIN vector. The INF2-ER[K792A] GFP construct was generated via site-directed mutagenesis using the QuikChange II Site-Directed Mutagenesis Kit (Agilent Technologies). All constructs were sequenced completely across their coding region.

### Data quantification and statistics

All graphs were generated using GraphPad Prism 10 software. All bar graphs display bars marking average values per condition and dots marking average values per cell analyzed. For all bar graphs, error bars indicate standard deviation. For all line graphs, thicker horizontal curves mark average values per condition and thin vertical lines indicate standard error of the mean. For all statistical comparisons, ^****^ indicates p-values≤0.0001, ^***^ indicates p-values≤0.001, ^**^ indicates p-values≤0.01, ^*^ indicates p-values≤0.05, and ns indicates p-values>0.05. All t-test and ANOVA statistical analyses were done using GraphPad Prism 10 software. Fisher’s exact test for comparison of observed versus expected values and the modified Chi-squared method for comparing curves were calculated using Microsoft Excel. The modified Chi-squared method has been previously described^30^. All line scans from Figure 1A and Supplementary Figure 1A-C were normalized and plotted in Microsoft Excel.

#### Organelle fission scoring

Organelle fission events were scored manually by visualizing the organelle channel in Fiji^31^ for each 5-minute time-lapse. To reduce the possibility of false positives caused by organelle crossover events or “kiss and run” events, fission events were classified as instances in which the organelle exists as a single organelle for at least 30 seconds prior to splitting into two or more organelles, which then remain separate for at least 30 seconds. For mitochondria, the presence of DRP1-mCherry at the site prior to fission was also checked. For quantifying presence or absence of AC-ER GFP at fission events, scorers were blinded to the AC-ER channel until after identification of the fission events.

#### Area of overlap and “by chance” estimations

Masks of AC-ER channels and organelle channels were generated in Fiji. For AC-ER, thresholding was set to mask only the top 25% of signal based on maximum pixel intensity. These settings match the thresholding settings used to visualize AC-ER when scoring its presence or absence at organelle fission events. All other masks were generated using default thresholding settings in Fiji, which uses the IsoData algorithm^32^ The integrated density of each mask was calculated. Areas of overlap between masks were generated using the image calculator tool in Fiji, and the integrated density of these areas was also measured. These values were used to calculate the percentage overlap (that is, integrated density for area of overlap between AC-ER and the organelle divided by integrated density for the organelle area yields the percentage of organelle area overlapped by AC-ER). This percentage was interpreted as the probability of AC-ER localizing to fission sites by chance. An example of this method is shown in Supplementary Figure 1D.

#### Quantification of organelle morphology

Fiji was used to quantify all readouts of organelle morphology. All analysis related to mitochondrial morphology (area, length, branching) was performed using the “Mitochondria Analyzer” plugin which uses a combination of image pre-processing, thresholding, particle analysis, and skeletonization to provide quantitative readouts of mitochondrial morphology^33^. Identical settings were used for all mitochondria images. For endosome and lysosome morphology measurements, a custom Fiji macro was written and applied identically to all images. This macro relies on the “MorphoLibJ” Fiji plugin^34^. The macro first pre-processed the image via Fiji’s “Subtract Background”, “Despeckle”, “Enhance Contrast”, and “Median Filter” tools to improve organelle signal and reduce noise. The image was then thresholded using Fiji’s default “Moments” algorithm^35^ to create a binary mask. The “Analyze Particles” function was then used to return a mask excluding all objects with a size less than 0.06 to further remove noise. To split touching objects, MorphoLibJ’s “Marker-controlled Watershed” tool was then used. Fiji’s “Find Maxima” function was used on the original (prior to binarization) image to create a point output which was used as the markers. Finally, MorphoLibJ’s “Analyze Regions” function was used to quantify the size and shape of each segmented object.

#### Quantification of organelle mobility via Imaris

For each organelle, all conditions were batch processed using identical settings. First, surfaces were created to segment individual organelles using Imaris’ Local Contrast threshold option. For tracking, Imaris’ autoregressive motion algorithm was used. Tracks with a duration less than 10 seconds were considered noise and filtered out. The “Track Displacement Distance” and “Track Duration” values calculated by Imaris were matched on a per organelle basis and used to calculate the “Mean Straight Velocity” (Track Displacement Distance divided by Track Duration) for each organelle. This calculation was done in Microsoft Excel.

#### Quantification of organelle mobility via autocorrelation analysis

As a supplement to the object tracking method used via Imaris software, we sought an additional method of measuring organelle movement which does not rely on organelle segmentation and can be performed using freely available software. Forgoing individual organelle segmentation eliminates most of the parameters that need to be carefully optimized for accurate object tracking. This simplifies analysis and may be more appropriate for measuring movement of organelles with complex morphologies, with the trade-off of missing potential details such as subpopulations and directionality. To this end, we wrote a custom Fiji macro which measures the correlation between the first frame of a time-lapse and all consecutive frames of the same time-lapse. The correlation values were measured using Fiji’s “Colocalization Threshold” function. Because correlation is a measure of the similarity between two images, the more organelles move, the more their positions differ compared to the first frame of the time-lapse, the less similar consecutive frames will be compared to the first frame, and the lower the correlation values will be. Put simply, the correlation values over time are inversely proportional to the degree of organelle movement. Thus, an increase in organelle movement should result in a faster decrease in the correlation coefficient over time and vice versa for a decrease in organelle movement. The macro saved a .csv file with the correlation values from each frame listed from the beginning to the end of the time-lapse. These values were then plotted to generate a curve of the decrease of the correlation values over time. The “Colocalization Threshold” function returns multiple types of correlation values. Here, we made comparisons using Manders’ Correlation Coefficient since, compared to the other readouts, it is less likely to be altered by factors unrelated to organelle movement such as bleaching of the organelle signal over time.

## Acknowledgements

The authors gratefully acknowledge Richard A. Kahn and members of the Shadel and Manor labs for feedback and suggestions throughout the project. This work was supported by NIH R01 AR069876, and the Allen-American Heart Association Initiative in Brain Health and Cognitive Impairment Award 19PABH134610000H, and the San Diego Nathan Shock Center P30AG068635 to G.S.S., who also holds the Audrey Geisel Chair in Biomedical Science. U.M. is supported by a CZI Imaging Scientist Award (DOI:10.37921/694870itnyzk) from the Chan Zuckerberg Initiative DAF), NSF NeuroNex Award 2014862, the UCSD Goeddel Family Technology Sandbox, and the Charcot-Marie-Tooth Association. This work was also supported by NIH 1F32GM137580 to C.R.S. Microscopy in this work was supported by the Waitt Advanced Biophotonics Core Facility of the Salk Institute for Biological Studies with funding from NIH-NCI CCSG: P30 014195 and the Waitt Foundation; the Transgenic and GT3 Core Facilities of the Salk Institute were supported with funding from NIH-NCI CCSG: P30 014195, an NINDS R24 Core Grant, and funding from NEI.

## Figures & Legends

**Supp Fig 1:**
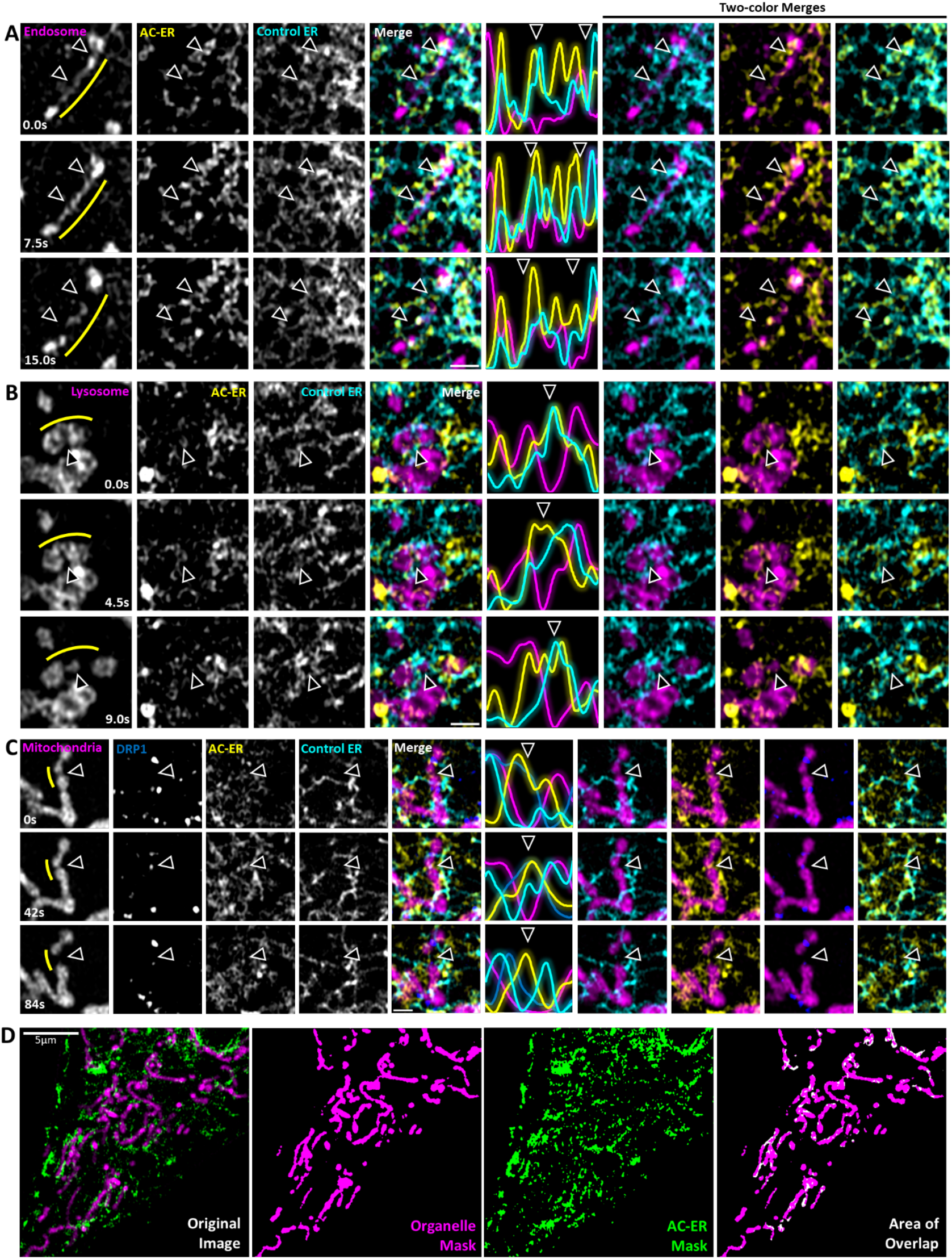
Organelle fission examples displaying all channels and example of masks used to generate “by chance” values. **A**. The same endosome example from Figure 1A is shown. Like Figure 1, arrowheads denote fission sites, and the yellow line denotes the location where the line scan was drawn. In order from left to right are single channel grayscale images, the merge of all channels (endosomes in magenta, AC-ER in yellow, and ER in cyan), the plot of pixel intensities corresponding to the line scan, and two-channel merges. Scale bar is 1µm. **B**. Same as A but displaying the lysosome example from Figure 1A in the magenta channel. **C**. Similar to A and B but displaying the mitochondria example from Figure 1A in the magenta channel. Additionally, the DRP1 channel is shown in blue. **D**. In the leftmost panel, an example of a typical cell used for analysis of area of overlap between organelles and AC-ER is shown. The organelles used in this example are mitochondria. Mitochondria are labeled in magenta and AC-ER is labeled in green. The appearance of the masks generated by the mitochondrial and AC-ER signals are shown in the center panels. The rightmost panel shows the area of overlap between the mitochondrial mask and AC-ER mask (white), overlaid with the mitochondrial mask (magenta). This area of overlap was used to determine the “by chance” values shown in Figure 1B and C.

**Supp Fig 2:**
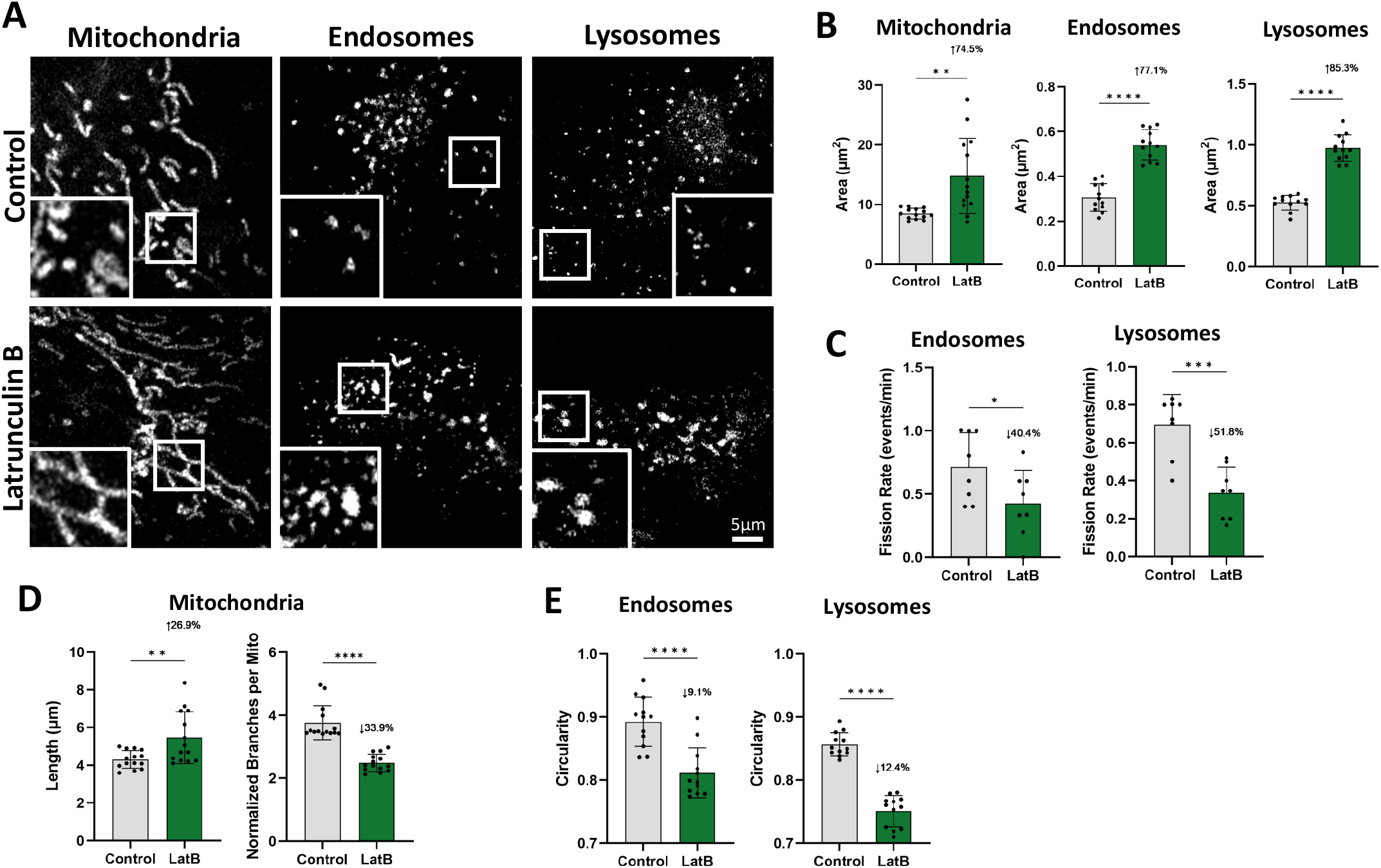
Inhibition of actin polymerization causes organelle enlargement. Organelles in U2OS cells were labeled with Rab5-mCherry (endosomes), LAMP1-mCherry (lysosomes), or MitoTracker Deep Red (mitochondria) and imaged live following addition of 200nM LatB or vehicle control for 60-90 minutes. **A**. Representative images of mitochondria, endosomes, and lysosomes after treatment with vehicle control or LatB are shown. Insets show magnified views of the boxed regions. **B**. Quantification of individual organelle area following treatment with LatB or vehicle control. Mitochondrial morphology was measured using the “Mitochondria Analyzer” Fiji plugin^33^. Endosome and lysosome morphology were measured using a custom Fiji macro which uses a combination of thresholding and marker-controlled watershed to split clustered organelles by using local maxima as seed points, allowing for segmentation of individual organelles. The areas of each individual organelle were measured and the average per cell is shown as black dots on the graphs. Bars show the average across all cells for each condition. Standard deviation is denoted by error bars. Conditions were compared via Welch’s t-test. N = 14 cells per condition for mitochondria and 12 per condition for endosomes and lysosomes. **C**. Quantification of manually scored fission rates in endosomes and lysosomes following LatB or vehicle control treatment. N = 8 cells per condition. **D**. Quantification of mitochondrial length and number of mitochondrial branches following LatB or vehicle control treatment. N = 14 cells per condition. **E**. Quantification of endosome and lysosome circularity following LatB or vehicle control treatment. N = 12 cells per condition. For all graphs, the magnitude of the change compared to control is shown. An up arrow indicates an increase and a down arrow indicates a decrease. For statistical comparisons ^****^ indicates p-value≤0.0001, ^***^ indicates p-values≤0.001, ^**^ indicates p-value≤0.01, and ^*^ indicates p-value≤0.05. All conditions were compared via Welch’s t-test. All experiments were performed with N=3 biological replicates.

**Supp Fig 3:**
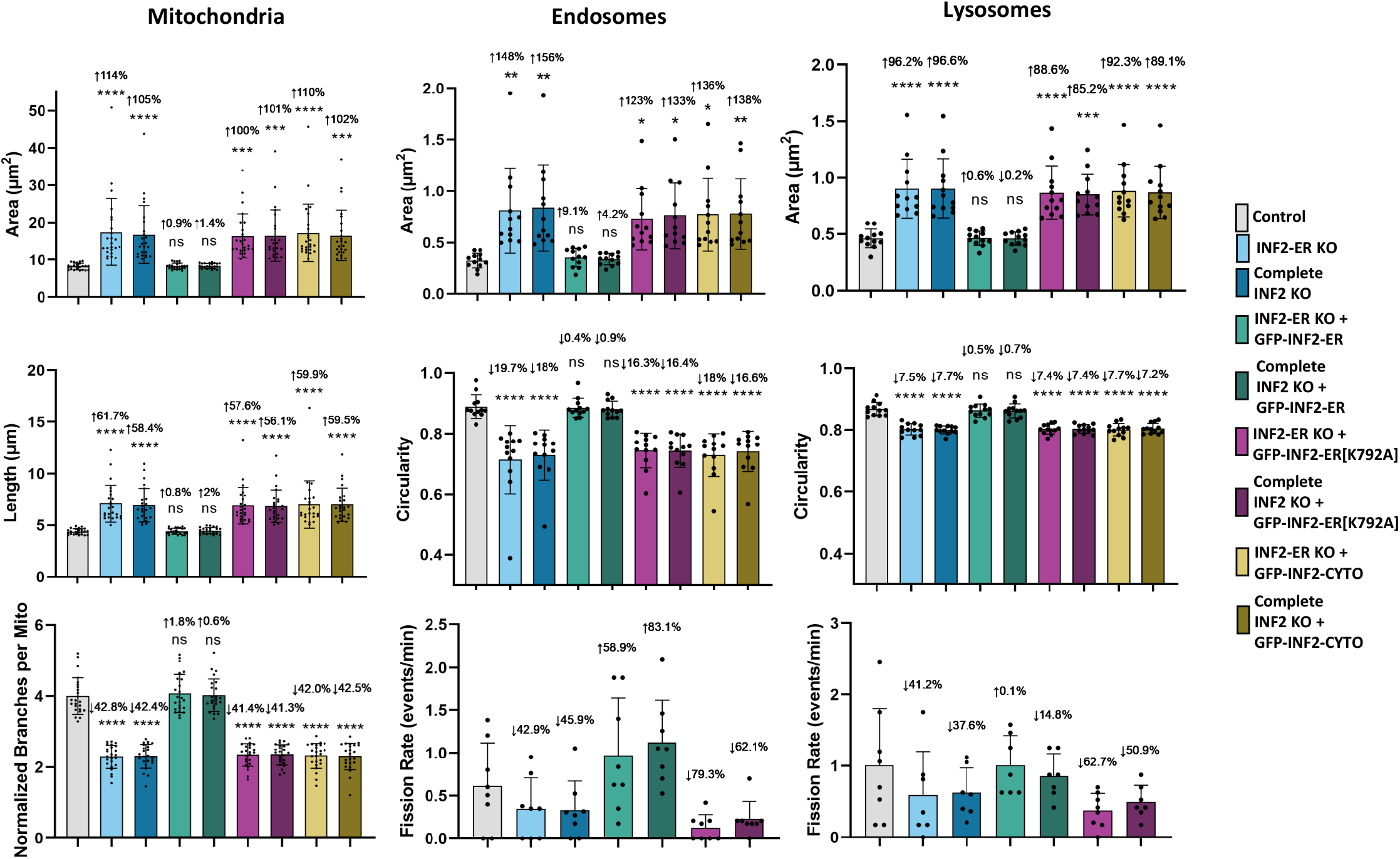
Further quantification of the effect of INF2 knock-out on organelle morphology. Additional quantification results related to the experiments described in Figure 2 are shown. The top row of graphs shows the same individual organelle area quantification results from Figure 2 with the addition of results from experiments carried out with U2OS cells where all isoforms of INF2 have been knocked-out (“Complete INF2 KO”). In addition to area, individual mitochondrial length, the number of branches per mitochondria, and circularity of endosomes and lysosomes are also shown. Endosome and lysosome fission rates are shown in the bottom right graphs. N = 8 cells per condition for fission rate results. For all other results, N = 12 cells per condition for endosomes and lysosomes and 24 cells per condition for mitochondria. For all graphs, black dots show the average per cell and bars indicate the average per condition. Error bars show standard deviation. The magnitude of the change and p-value compared to control is shown. An up arrow indicates an increase and a down arrow indicates a decrease. For statistical comparisons ^****^ indicates p-value≤0.0001, ^***^ indicates p-values≤0.001, ^**^ indicates p-value≤0.01, ^*^ indicates p-value≤0.05, and ns indicates p-value>0.05. Conditions were compared via ordinary one-way ANOVA. All experiments were performed with N=3 biological replicates.

**Supp Fig 4:**
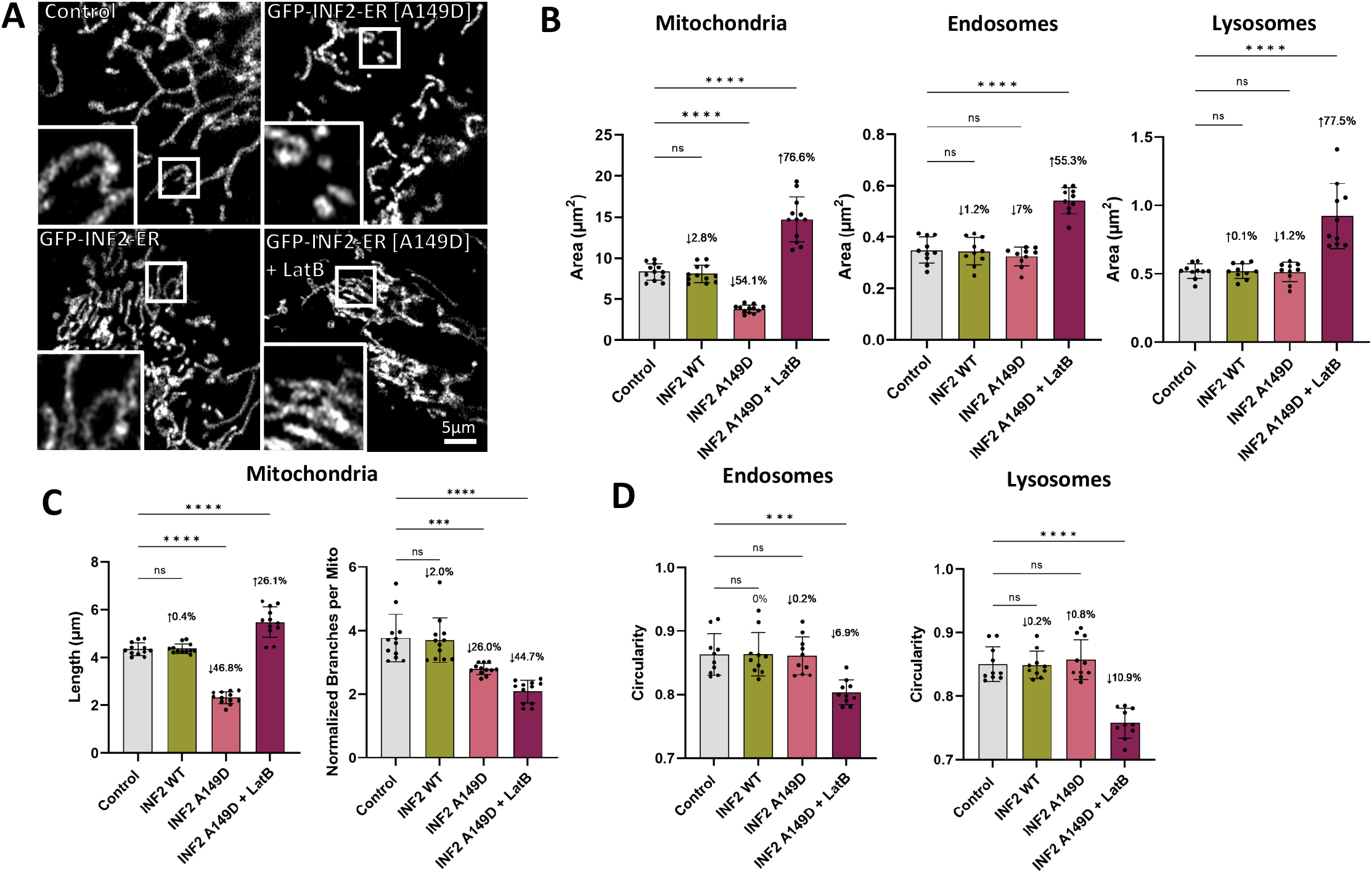
Expression of dominant active INF2 causes mitochondrial fragmentation but does not alter endosome or lysosome shape. Wild-type U2OS cells were imaged live following co-transfection with organelle labels and various INF2-ER constructs. **A**. Representative images of mitochondria labeled with MitoTracker Deep Red in wild-type U2OS cells expressing GFP (control), wild-type INF2-ER GFP, or INF2-ER[A149D] GFP are shown. A representative image of mitochondria from the INF2-ER[A149D] GFP condition following treatment with 1µM LatB for 15-30 minutes is also shown. **B**. Quantification of individual organelle area for each condition and organelle. **C**. Quantification of mitochondrial length and branching for each condition. **D**. Quantification of endosome and lysosome circularity for all conditions. For all graphs, black dots show the average per cell and bars indicate the average per condition. Error bars show standard deviation. The magnitude of the change and p-value compared to control is shown. An up arrow indicates an increase and a down arrow indicates a decrease. For statistical comparisons ^****^ indicates p-value≤0.0001, ^***^ indicates p-values≤0.001, ^**^ indicates p-value≤0.01, ^*^ indicates p-value≤0.05, and ns indicates p-value>0.05. Conditions were compared via ordinary one-way ANOVA. N = 10 cells per condition for endosomes and lysosomes and 12 cells per condition for mitochondria. All experiments were performed with N=3 biological replicates.

**Fig S5.**
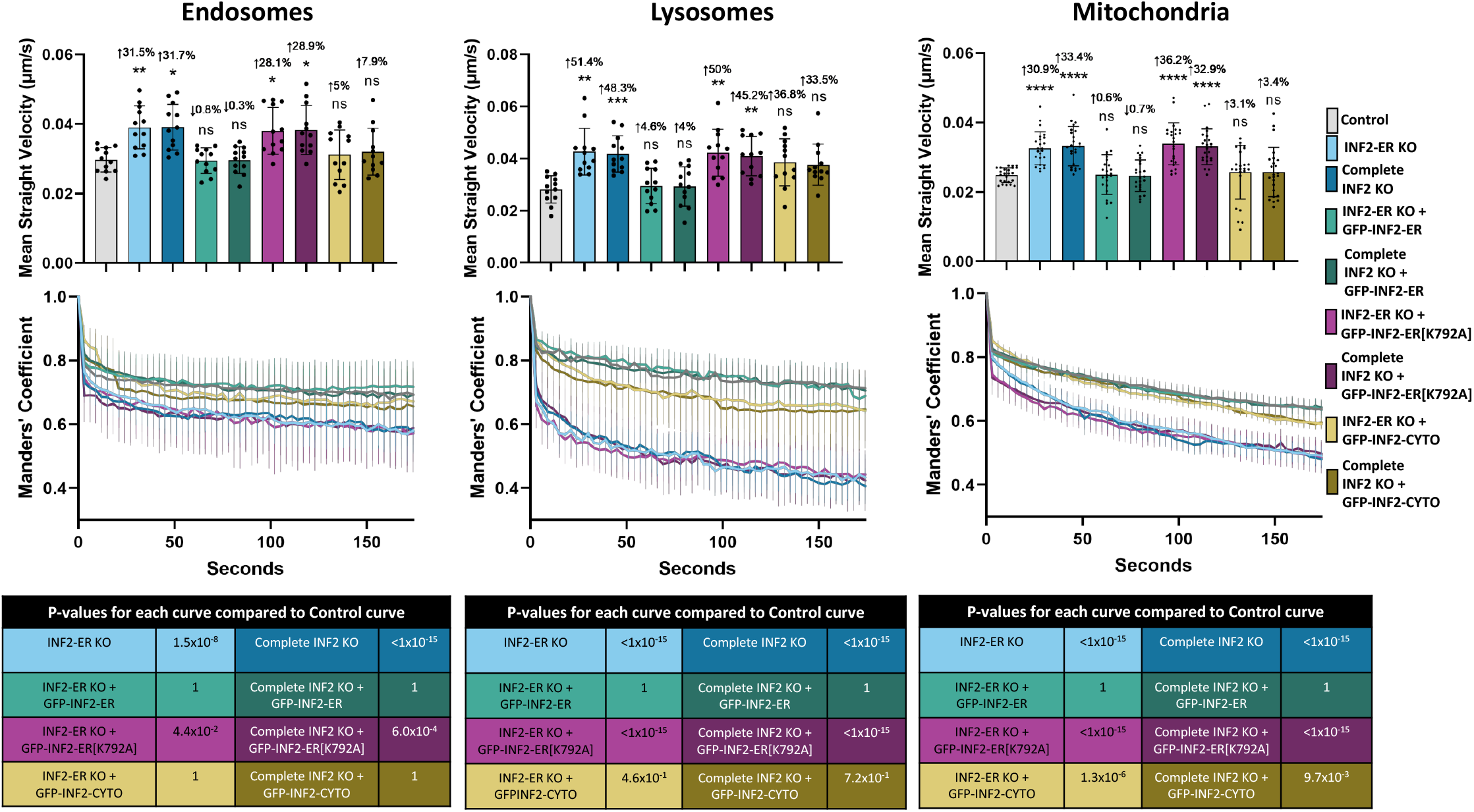
Supp Fig 5: Further quantification of the effect of INF2 KO on organelle mobility. Additional quantification results related to the experiments described in Figure 3 are shown. The top row of graphs shows the same mean straight velocity values from Figure 3 with the addition of values from experiments carried out in Complete INF2 KO cells. Black dots show the average per cell and bars indicate the average per condition. Error bars show standard deviation. The magnitude of the change and p-value compared to control is shown. An up arrow indicates an increase and a down arrow indicates a decrease. For statistical comparisons ^****^ indicates p-value≤0.0001, ^***^ indicates p-values≤0.001, ^**^ indicates p-value≤0.01, ^*^ indicates p-value≤0.05, and ns indicates p-value>0.05. Conditions were compared via ordinary one-way ANOVA. The second row of graphs shows the autocorrelation values of the organelle channel over time for each condition calculated using a custom Fiji macro described in Methods. The thick lines mark the average correlation value over time and the thin vertical lines show the standard error. Tables below the graphs show the p-value results of comparing each curve to the control curve. Curves were compared using a modified Chi-squared method^30^. For all graphs, N = 12 cells per condition for endosomes and lysosomes and 24 cells per condition for mitochondria. All experiments were performed with N=3 biological replicates.

**Supp Fig 6:**
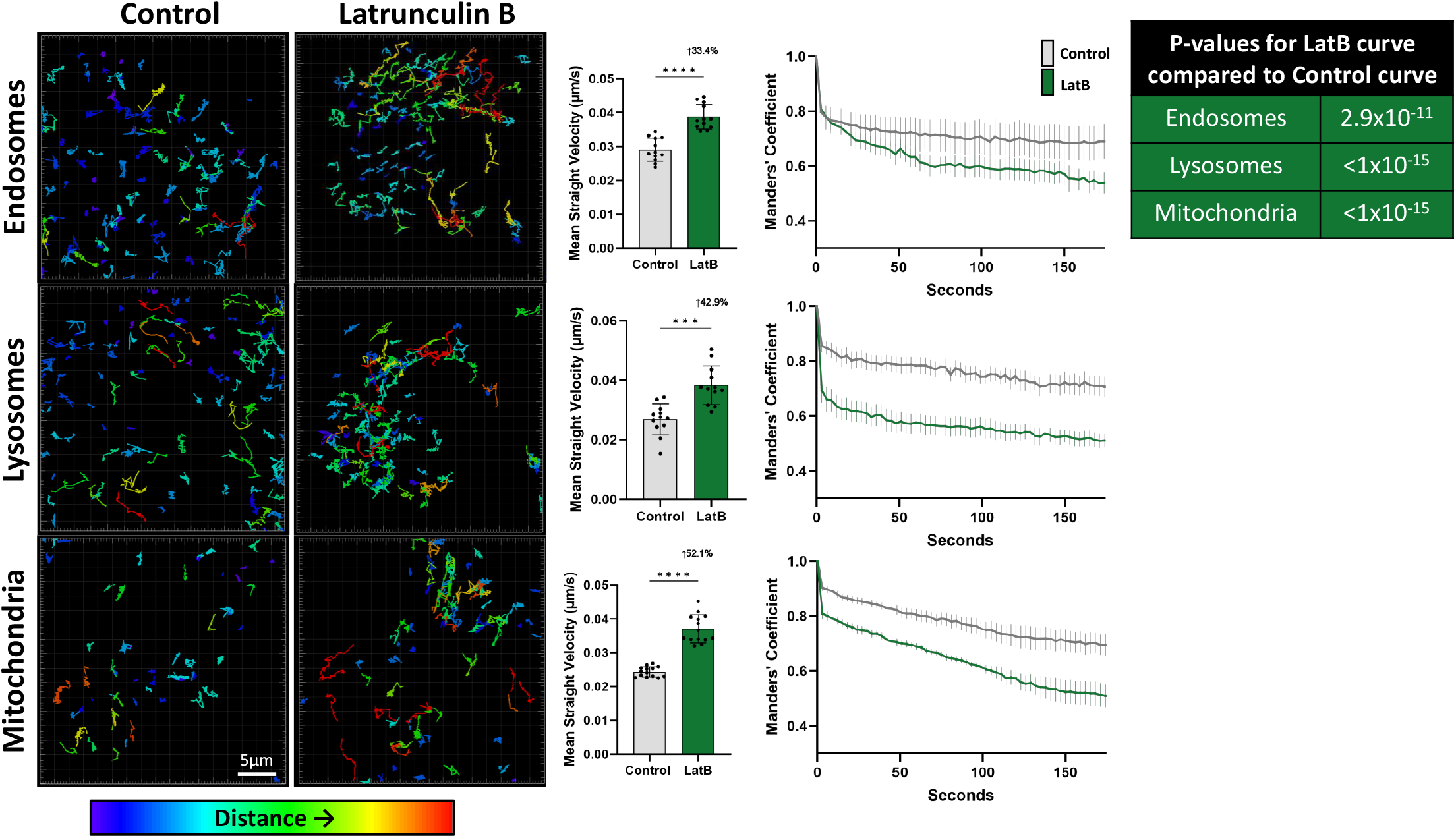
Inhibition of actin polymerization increases organelle mobility. Wild-type U2OS cells were labeled with Rab5-mCherry (endosomes), LAMP1-mCherry (lysosomes), or MitoTracker Deep Red (mitochondria), treated with 200nM LatB or vehicle control for 60-90 minutes, then imaged over time for an additional 5 minutes. Organelle movement was measured using both Imaris tracking and our autocorrelation method. Panels display tracks of individual organelle trajectories color-coded based on distance traveled as described in Figure 3. Graphs of organelle tracking results are also displayed. Black dots show the average per cell and bars indicate the average per condition. Error bars show standard deviation. The magnitude of the change and p-value compared to control is shown. An up arrow indicates an increase and a down arrow indicates a decrease. For statistical comparisons ^****^ indicates p-value≤0.0001, ^***^ indicates p-values≤0.001, ^**^ indicates p-value≤0.01, ^*^ indicates p-value≤0.05, and ns indicates p-value>0.05. Conditions were compared via Welch’s t-test. Graphs of the autocorrelation values over time are also shown. The thick lines mark the average correlation value over time and the thin vertical lines show the standard error. The table displays the p-values resulting from comparing the control and LatB curves. For all graphs, N = 12 cells per condition for endosomes and lysosomes and 14 cells per condition for mitochondria. All experiments were performed with N=3 biological replicates.

**Supp Fig 7:**
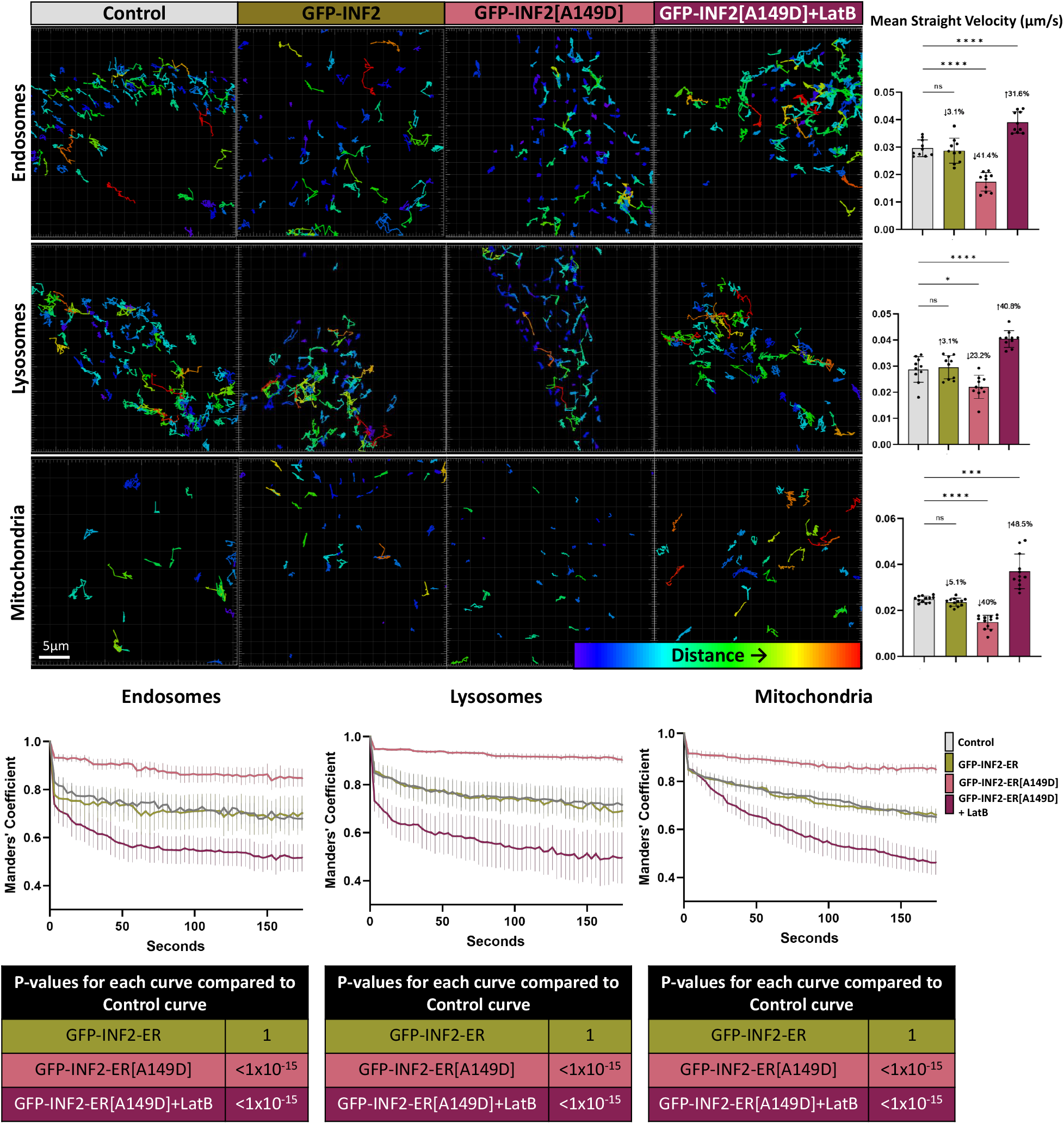
Expression of dominant active INF2 reduces organelle mobility. In wild-type U2OS cells organelles were labeled using Rab5-mCherry (endosomes), LAMP1-mCherry (lysosomes), or MitoTracker Deep Red (mitochondria) and co-transfected with GFP (control), wild-type INF2-ER GFP, or INF2-ER[A149D] GFP. The INF2-ER[A149D] GFP condition was also compared with and without 15-30 minute treatment with 1µM LatB. Tracks displaying individual organelle trajectories color-coded by distance traveled are shown for representative cells from each condition. Quantification of these results via organelle tracking in Imaris is shown in the bar graphs. For all bar graphs, black dots show the average per cell and bars indicate the average per condition. Error bars show standard deviation. The magnitude of the change and p-value compared to control are shown. An up arrow indicates an increase and a down arrow indicates a decrease. For statistical comparisons, ^****^ indicates p-value≤0.0001, ^***^ indicates p-values≤0.001, ^**^ indicates p-value≤0.01, ^*^ indicates p-value≤0.05, and ns indicates p-value>0.05. Conditions were compared via ordinary one-way ANOVA. In addition to object tracking in Imaris, organelle movement was also quantified using our autocorrelation method, displayed in the line graphs. The thick lines mark the average correlation value over time and the thin vertical lines show the standard error. The tables display the p-values resulting from comparing each curve with the control curve. N = 10 cells per condition for endosomes and lysosomes and 12 cells per condition for mitochondria. All experiments were performed with N=3 biological replicates.

**Supp Video 1: ER-associated actin accumulates at endosomal fission sites prior to endosome division**. Timelapse video corresponding to the endosome example from Figure 1A and Supplemental Figure 1A. The top row shows single channel grayscale videos of the endosome (Rab5-mCherry), AC-ER (AC-ER GFP), and ER (BFP-KDEL) channels followed by a merge of all channels with endosomes in magenta, AC-ER in yellow, and ER in cyan. The bottom row shows two-color merges following the same color-scheme. White arrowheads indicate the location of endosome fission events. Scale bar = 1µm.

**Supp Video 2: ER-associated actin accumulates at lysosomal fission sites prior to lysosome division**. Timelapse video corresponding to the lysosome example from Figure 1A and Supplemental Figure 1B. The top row shows single channel grayscale videos of the lysosome (LAMP1-mCherry), AC-ER (AC-ER GFP), and ER (BFP-KDEL) channels followed by a merge of all channels with lysosomes in magenta, AC-ER in yellow, and ER in cyan. The bottom row shows two-color merges following the same color-scheme. White arrowheads indicate the location of lysosome fission events. Scale bar = 1µm.

**Supp Video 3: ER-associated actin accumulates at mitochondrial fission sites prior to mitochondrial division**. Timelapse video corresponding to the mitochondria example from Figure 1A and Supplemental Figure 1C. The top row shows single channel grayscale videos of the mitochondria (MitoTracker Deep Red), DRP1 (DRP1-mCherry), AC-ER (AC-ER GFP), and ER (BFP-KDEL) channels followed by a merge of all channels with mitochondria in magenta, DRP1 in blue, AC-ER in yellow, and ER in cyan. The bottom row shows two-color merges following the same color-scheme. White arrowheads indicate the location of mitochondrial fission events. Scale bar = 1µm.

**Supp Video 4: Loss of INF2 increases endosome mobility**. Timelapse video corresponding to the endosome examples from Figure 3. The endosome channel (Rab5-mCherry) is shown for the indicated conditions. Scale bar = 10µm.

**Supp Video 5: Loss of INF2 increases lysosome mobility**. Timelapse video corresponding to the lysosome examples from Figure 3. The lysosome channel (LAMP1-mCherry) is shown for the indicated conditions. Scale bar = 10µm.

**Supp Video 6: Loss of INF2 increases mitochondrial mobility**. Timelapse video corresponding to the mitochondria examples from Figure 3. The mitochondria channel (MitoTracker Deep Red) is shown for the indicated conditions. Scale bar = 10µm.

**Supp Video 7: Inhibition of actin polymerization increases organelle mobility**. Timelapse video corresponding to Supplementary Figure 6. The first column shows the organelle channel from U2OS cells treated with vehicle control and the second column shows the organelle channel from U2OS cells treated with 200nM LatB for 60-90 minutes prior to timelapse acquisition. The top row shows endosomes (Rab5-mCherry), the middle row shows lysosomes (LAMP1-mCherry), and the bottom row shows mitochondria (MitoTracker Deep Red). Scale bar = 10µm.

**Supp Video 8: Expression of dominant active INF2 reduces endosome mobility**. Timelapse video corresponding to the endosome example from Supplementary Figure 7. The endosome channel (Rab5-mCherry) is shown for the indicated conditions. The LatB-treated condition (+LatB) was treated with 1µM LatB for 15-30 minutes prior to timelapse acquisition. Scale bar = 10µm.

**Supp Video 9: Expression of dominant active INF2 reduces lysosome mobility**. Timelapse video corresponding to the lysosome example from Supplementary Figure 7. The lysosome channel (LAMP1-mCherry) is shown for the indicated conditions. The LatB-treated condition (+LatB) was treated with 1µM LatB for 15-30 minutes prior to timelapse acquisition. Scale bar = 10µm.

**Supp Video 10: Expression of dominant active INF2 reduces mitochondrial mobility**. Timelapse video corresponding to the mitochondria example from Supplementary Figure 7. The mitochondria channel (MitoTracker Deep Red) is shown for the indicated conditions. The LatB-treated condition (+LatB) was treated with 1µM LatB for 15-30 minutes prior to timelapse acquisition. Scale bar = 10µm.

## References

1. Krols M, van Isterdael G, Asselbergh B, Kremer A, Lippens S, Timmerman V, Janssens S. Mitochondria-associated membranes as hubs for neurodegeneration. Acta neuropathologica. 2016;131(4):505–23 doi: 10.1007/s00401-015-1528-7. PubMed PMID: 26744348; PMCID: 4789254.

2. Paillusson S, Stoica R, Gomez-Suaga P, Lau DHW, Mueller S, Miller T, Miller CCJ. There’s Something Wrong with my MAM; the ER-Mitochondria Axis and Neurodegenerative Diseases. Trends in neurosciences. 2016;39(3):146–57. doi: 10.1016/j.tins.2016.01.008. PubMed PMID: 26899735; PMCID: 4780428.

3. Arasaki K, Shimizu H, Mogari H, Nishida N, Hirota N, Furuno A, Kudo Y, Baba M, Baba N, Cheng J, Fujimoto T, Ishihara N, Ortiz-Sandoval C, Barlow LD, Raturi A, Dohmae N, Wakana Y, Inoue H, Tani K, Dacks JB, Simmen T, Tagaya M. A role for the ancient SNARE syntaxin 17 in regulating mitochondrial division. Developmental cell. 2015;32(3):304–17. Epub 2015/01/27. doi: 10.1016/j.devcel.2014.12.011. PubMed PMID: 25619926.

4. Area-Gomez E, de Groof AJ, Boldogh I, Bird TD, Gibson GE, Koehler CM, Yu WH, Duff KE, Yaffe MP, Pon LA, Schon EA. Presenilins are enriched in endoplasmic reticulum membranes associated with mitochondria. The American journal of pathology. 2009;175(5):1810–6. Epub 2009/10/17. doi: 10.2353/ajpath.2009.090219. PubMed PMID: 19834068; PMCID: Pmc2774047.

5. Area-Gomez E, Del Carmen Lara Castillo M, Tambini MD, Guardia-Laguarta C, de Groof AJ, Madra M, Ikenouchi J, Umeda M, Bird TD, Sturley SL, Schon EA. Upregulated function of mitochondria-associated ER membranes in Alzheimer disease. The EMBO journal. 2012;31(21):4106–23. Epub 2012/08/16. doi: 10.1038/emboj.2012.202. PubMed PMID: 22892566; PMCID: Pmc3492725.

6. Schon EA, Przedborski S. Mitochondria: the next (neurode)generation. Neuron. 2011;70(6):1033–53. Epub 2011/06/22. doi: 10.1016/j.neuron.2011.06.003. PubMed PMID: 21689593; PMCID: Pmc3407575.

7. Stoica R, De Vos KJ, Paillusson S, Mueller S, Sancho RM, Lau KF, Vizcay-Barrena G, Lin WL, Xu YF, Lewis J, Dickson DW, Petrucelli L, Mitchell JC, Shaw CE, Miller CC. ER-mitochondria associations are regulated by the VAPB-PTPIP51 interaction and are disrupted by ALS/FTD-associated TDP-43. Nature communications. 2014;5:3996. Epub 2014/06/04. doi: 10.1038/ncomms4996. PubMed PMID: 24893131; PMCID: Pmc4046113.

8. Gong G, Song M, Csordas G, Kelly DP, Matkovich SJ, Dorn GW, 2nd. Parkin-mediated mitophagy directs perinatal cardiac metabolic maturation in mice. Science. 2015;350(6265):aad2459. Epub 2016/01/20. doi: 10.1126/science.aad2459. PubMed PMID: 26785495; PMCID: PMC4747105.

9. Rowland AA, Voeltz GK. Endoplasmic reticulum-mitochondria contacts: function of the junction. Nat Rev Mol Cell Biol. 2012;13(10):607–25. doi: 10.1038/nrm3440. PubMed PMID: 22992592; PMCID: 5111635.

10. Friedman JR, Lackner LL, West M, DiBenedetto JR, Nunnari J, Voeltz GK. ER tubules mark sites of mitochondrial division. Science. 2011;334(6054):358–62. Epub 2011/09/03. doi: 10.1126/science.1207385. PubMed PMID: 21885730; PMCID: PMC3366560.

11. Lewis SC, Uchiyama LF, Nunnari J. ER-mitochondria contacts couple mtDNA synthesis with mitochondrial division in human cells. Science. 2016;353(6296). doi: 10.1126/science.aaf5549.

12. Lee JE, Cathey PI, Wu H, Parker R, Voeltz GK. Endoplasmic reticulum contact sites regulate the dynamics of membraneless organelles. Science. 2020;367(6477). doi: 10.1126/science.aay7108. PubMed PMID: 32001628; PMCID: PMC10088059.

13. Hoyer MJ, Chitwood PJ, Ebmeier CC, Striepen JF, Qi RZ, Old WM, Voeltz GK. A Novel Class of ER Membrane Proteins Regulates ER-Associated Endosome Fission. Cell. 2018;175(1):254–65 e14. Epub 20180913. doi: 10.1016/j.cell.2018.08.030. PubMed PMID: 30220460; PMCID: PMC6195207.

14. Hatch AL, Ji WK, Merrill RA, Strack S, Higgs HN. Actin filaments as dynamic reservoirs for Drp1 recruitment. Mol Biol Cell. 2016;27(20):3109–21. Epub 2016/08/26. doi: 10.1091/mbc.E16-03-0193. PubMed PMID: 27559132; PMCID: PMC5063618.

15. Korobova F, Ramabhadran V, Higgs HN. An actin-dependent step in mitochondrial fission mediated by the ER-associated formin INF2. Science. 2013;339(6118):464–7. doi: 10.1126/science.1228360. PubMed PMID: 23349293; PMCID: 3843506.

16. Korobova F, Gauvin TJ, Higgs HN. A role for myosin II in mammalian mitochondrial fission. Curr Biol. 2014;24(4):409–14. Epub 2014/02/04. doi: 10.1016/j.cub.2013.12.032. PubMed PMID: 24485837; PMCID: PMC3958938.

17. Saffi GT, Botelho RJ. Lysosome Fission: Planning for an Exit. Trends Cell Biol. 2019;29(8):635–46. Epub 20190603. doi: 10.1016/j.tcb.2019.05.003. PubMed PMID: 31171420.

18. Gautreau A, Oguievetskaia K, Ungermann C. Function and regulation of the endosomal fusion and fission machineries. Cold Spring Harb Perspect Biol. 2014;6(3). Epub 20140301. doi: 10.1101/cshperspect.a016832. PubMed PMID: 24591520; PMCID: PMC3949357.

19. Schiavon CR, Zhang T, Zhao B, Moore AS, Wales P, Andrade LR, Wu M, Sung T-C, Dayn Y, Feng JW, Quintero OA, Shadel GS, Grosse R, Manor U. Actin chromobody imaging reveals sub-organellar actin dynamics. Nature Methods. 2020;17(9):917–21. doi: 10.1038/s41592-020-0926-5.

20. De Vos KJ, Allan VJ, Grierson AJ, Sheetz MP. Mitochondrial function and actin regulate dynamin-related protein 1-dependent mitochondrial fission. Curr Biol. 2005;15(7):678–83. doi: 10.1016/j.cub.2005.02.064. PubMed PMID: 15823542.

21. Hatch AL, Gurel PS, Higgs HN. Novel roles for actin in mitochondrial fission. J Cell Sci. 2014;127(Pt 21):4549–60. Epub 2014/09/14. doi: 10.1242/jcs.153791. PubMed PMID: 25217628; PMCID: PMC4215709.

22. Li S, Xu S, Roelofs BA, Boyman L, Lederer WJ, Sesaki H, Karbowski M. Transient assembly of F-actin on the outer mitochondrial membrane contributes to mitochondrial fission. The Journal of cell biology. 2015;208(1):109–23. Epub 20141229. doi: 10.1083/jcb.201404050. PubMed PMID: 25547155; PMCID: PMC4284235.

23. Ramabhadran V, Gurel PS, Higgs HN. Mutations to the formin homology 2 domain of INF2 protein have unexpected effects on actin polymerization and severing. J Biol Chem. 2012;287(41):34234–45. Epub 2012/08/11. doi: 10.1074/jbc.M112.365122. PubMed PMID: 22879592; PMCID: PMC3464531.

24. Giardini PA, Theriot JA. Effects of intermediate filaments on actin-based motility of Listeria monocytogenes. Biophys J. 2001;81(6):3193–203. doi: 10.1016/S0006-3495(01)75955-3. PubMed PMID: 11720985; PMCID: PMC1301779.

25. Ji WK, Hatch AL, Merrill RA, Strack S, Higgs HN. Actin filaments target the oligomeric maturation of the dynamin GTPase Drp1 to mitochondrial fission sites. Elife. 2015;4:e11553. Epub 20151126. doi: 10.7554/eLife.11553. PubMed PMID: 26609810; PMCID: PMC4755738.

26. Liu A, Kage F, Higgs HN. Mff oligomerization is required for Drp1 activation and synergy with actin filaments during mitochondrial division. Mol Biol Cell. 2021;32(20):ar5. Epub 2021/08/05. doi: 10.1091/mbc.E21-04-0224. PubMed PMID: 34347505; PMCID: PMC8684745.

27. Schiavon CR, Shadel GS, Manor U. Impaired Mitochondrial Mobility in Charcot-Marie-Tooth Disease. Front Cell Dev Biol. 2021;9:624823. Epub 2021/02/19. doi: 10.3389/fcell.2021.624823. PubMed PMID: 33598463; PMCID: PMC7882694.

28. Chakrabarti R, Ji WK, Stan RV, de Juan Sanz J, Ryan TA, Higgs HN. INF2-mediated actin polymerization at the ER stimulates mitochondrial calcium uptake, inner membrane constriction, and division. The Journal of cell biology. 2018;217(1):251–68. doi: 10.1083/jcb.201709111. PubMed PMID: 29142021; PMCID: 5748994.

29. Mali P, Yang L, Esvelt KM, Aach J, Guell M, DiCarlo JE, Norville JE, Church GM. RNA-guided human genome engineering via Cas9. Science. 2013;339(6121):823–6. Epub 20130103. doi: 10.1126/science.1232033. PubMed PMID: 23287722; PMCID: PMC3712628.

30. Hristova K, Wimley WC. Determining the statistical significance of the difference between arbitrary curves: A spreadsheet method. PLoS One. 2023;18(10):e0289619. Epub 20231031. doi: 10.1371/journal.pone.0289619. PubMed PMID: 37906570; PMCID: PMC10617697.

31. Schindelin J, Arganda-Carreras I, Frise E, Kaynig V, Longair M, Pietzsch T, Preibisch S, Rueden C, Saalfeld S, Schmid B, Tinevez JY, White DJ, Hartenstein V, Eliceiri K, Tomancak P, Cardona A. Fiji: an open-source platform for biological-image analysis. Nat Methods. 2012;9(7):676–82. Epub 20120628. doi: 10.1038/nmeth.2019. PubMed PMID: 22743772; PMCID: PMC3855844.

32. Ridler TW, Calvard S. Picture Thresholding Using an Iterative Selection Method. Ieee T Syst Man Cyb. 1978;8(8):630-2. doi: DOI 10.1109/tsmc.1978.4310039. PubMed PMID: WOS:A1978FJ91600005.

33. Chaudhry A, Shi R, Luciani DS. A pipeline for multidimensional confocal analysis of mitochondrial morphology, function, and dynamics in pancreatic β-cells. Am J Physiol Endocrinol Metab. 2020;318(2):E87–e101. Epub 20191217. doi: 10.1152/ajpendo.00457.2019. PubMed PMID: 31846372; PMCID: PMC7052579.

34. Legland D, Arganda-Carreras I, Andrey P. MorphoLibJ: integrated library and plugins for mathematical morphology with ImageJ. Bioinformatics. 2016;32(22):3532–4. Epub 20160713. doi: 10.1093/bioinformatics/btw413. PubMed PMID: 27412086.

35. Tsai W-H. Moment-preserving thresolding: A new approach. Comput Vis Graph Image Process. 1985;29:377–93.

